# An annotated bioacoustic dataset of dog vocalizations and related sounds during dog-human social play

**DOI:** 10.64898/2026.04.20.719471

**Authors:** Laura V. Cuaya, Paula Pérez-Fraga, Raúl Hernández-Pérez, Louisa Pillwax, Ines Waldecker, Csenge Reisinger, Tamás Faragó, Sasha Winkler, Ludwig Huber, Claus Lamm

**Author notes:** Corresponding author: Laura V. Cuaya. These authors contributed equally (Ludwig Huber, Claus Lamm).

## Abstract

Dog (*Canis lupus familiaris*) vocal communication occurs across a wide range of social contexts, including dog–human social play, a common and dynamic interaction in which diverse vocal signals are expressed, particularly in young dogs. However, most available open datasets of dog vocalizations focus primarily on barks, leaving other sound types underrepresented. Here we present a bioacoustic dataset of dog–human social play sessions recorded under standardized laboratory conditions, comprising 30 play sessions involving 17 young dogs (6–24 months old) of different breeds playing with familiar humans. Raw audio recordings are accompanied by two layers of annotations covering the dog sounds produced during sessions. Video recordings of the sessions are provided for contextual reference, along with metadata describing each dog and the experimental sessions. Additionally, permutation-based classification analysis showed that annotated sound categories exhibit above-chance and generalizable acoustic differences across individuals. The dataset may support research on dog vocal communication and expand the range of sounds documented during positive dog–human social interactions.

## Background and Summary

Play is a widespread behaviour in dogs (*Canis lupus familiaris*) characterised by dynamic interactions such as chasing, play bows, physical contact, and frequently accompanied by vocal signals associated with arousal and social engagement, including barks, growls, grunts, and panting^1–7^. In addition to playing with conspecifics, domestic dogs readily engage in social play with humans, creating a distinctive form of interspecific interaction involving coordinated behaviours and shared engagement between partners^8,9^. Social play in dogs is influenced by factors such as age and partner familiarity: play occurs most frequently during juvenile stages and dogs typically engage more readily in play with familiar humans than with unfamiliar individuals^9,10^. These characteristics make dog–human play a naturalistic and particularly informative context for studying dog vocal communication in positive social settings.

Dogs possess a diverse vocal repertoire, and their vocal signals appear to encode both category-related information, linked to context and affective state, and individual-specific information, linked to caller morphology. For example, barks vary acoustically across contexts such as play and aggression-related contexts, whereas growls and whines also provide cues to caller body size^11–14^. However, this broader diversity of dog sounds remains only partly represented in currently available open datasets, particularly for dog-human social play.

Several open datasets of dog vocalizations have been developed to support research in animal communication, bioacoustics, and machine learning. For instance, DogSpeak^15^ consists of a large-scale collection of annotated bark sequences, while Dog2vec^16^ introduces a self-supervised representation learning framework trained on dog bark vocalizations. Other datasets address detection and classification tasks, including Barkopedia^17^, which supports automatic detection of dog barks, and BarkMeowDB^18^, a dataset frequently used in audio classification experiments involving dog barks and cat meows. Additional resources include the Dog Voice Emotion Dataset^19^, which contains labeled bark, growl, and grunt vocalizations; EmotionalCanines^20^, a dataset of dog bark clips annotated with corresponding arousal and valence labels; recordings of puppy whines produced during separation from the mother^21^; and AudioSet^22^, whose Dog category includes annotated audio clips of bark, howl, growl, and whimper vocalizations. While these datasets provide valuable resources for studying dog vocalizations and for developing automatic classification methods, most existing resources focus primarily on a limited subset of dog vocalization types, particularly barks. In addition, many datasets compile recordings from diverse sources such as online media or field recordings. Although this diversity is useful for training and evaluating computational models, it may also lead to a relative underrepresentation of more subtle sounds produced during social interactions, such as panting, which commonly occur during social play.

To our knowledge, no datasets document the full range of dog sounds produced during dog-human social play. To address this gap, we present a bioacoustic dataset consisting of 30 laboratory-recorded sessions of dog-human social play. The dataset includes raw audio recordings from seventeen dogs (6–24 months old) of different breeds interacting with a familiar human under standardized conditions. In addition to the audio recordings, the dataset provides two layers of annotations of dog sounds in the audio recordings, video recordings of the sessions for contextual reference, and metadata describing both the participating dogs and the experimental sessions. A limitation of the dataset is that it focuses on young dogs interacting with familiar humans in a specific type of play under controlled laboratory conditions, and therefore does not capture the full diversity of play contexts, including interactions with unfamiliar humans, different types of play activities, or play with conspecifics. Nevertheless, the dataset provides structured recordings of vocal behaviour during dog-human social play and may support research on canine vocal communication, behavioural interactions, and computational analysis of animal sounds.

## Methods

### Participants

Seventeen pet dogs were recorded for this study (Table 1). The sample included 10 males and 7 females, with an age range of 6 to 24 months (mean age = 14.94 months, SD = 6.36). The sample included a variety of breeds, specifically eight mixed-breed dogs, two Beaucerons, two Nova Scotia Duck Tolling Retrievers, one American Pit Bull Terrier, one Belgian Shepherd Tervuren, one Border Collie, one Fox Terrier, and one Mudi.

**Table 1.** Participant demographics and recording information.

| ID | Name | Breed | Sex | Weight<br>(kg) | Age<sup>1</sup> | Microphones |  |  | Raw<br>audios (s) | Views<sup>2</sup> |
| --- | --- | --- | --- | --- | --- | --- | --- | --- | --- | --- |
|  |  |  |  |  |  | M<sup>1</sup> | M<sup>2</sup> | M<sup>3</sup> |  |  |
| 01 | Simon | Mix | M | 40 | 12 | Y | <u>Y</u> | N | 390 | 7 |
| 02 | Lolka | Mix | M | 8.5 | 6 | Y | <u>Y</u> | Y | 552 | 7 |
| 03 | Falatka | Mix | F | 10 | 18 | Y | <u>Y</u> | Y | 512 | 7 |
| 04 | Mirabell | Mudi | F | 7 | 6 | Y | <u>Y</u> | Y | 689 | 7 |
| 05 | Koga | Mix | M | 18 | 15 | Y | N | <u>Y</u> | 690 | 7 |
| 06 | Csieko | Mix | F | 17 | 24 | Y | N | <u>Y</u> | 548 | 7 |
| 07 | Tofu | Fox terrier | F | 9 | 18 | Y | <u>Y</u> | Y | 846 | 7 |
| 08 | Aisha | BST | F | 15 | 10 | Y | <u>Y</u> | Y | 577 | 7 |
| 09 | CsiCsi | Beauceron | M | 32 | 12 | Y | <u>Y</u> | Y | 536 | 7 |
| 10 | Nitro | Beauceron | M | 45 | 24 | Y | <u>Y</u> | Y | 472 | 6 |
| 11 | Rudoph | NSCTR | M | 18 | 12 | Y | <u>Y</u> | Y | 465 | 7 |
| 12 | Kifli | Mix | M | 10 | 18 | Y | <u>Y</u> | Y | 497 | 7 |
| 13 | Rudy | Border collie | M | 17 | 12 | Y | <u>Y</u> | Y | 620 | 6 |
| 14 | Pigmy | NSCTR | F | 14.8 | 12 | Y | <u>Y</u> | Y | 519 | 6 |
| 15 | Bukta | Mix | F | 10 | 7 | Y | <u>Y</u> | Y | 1468 | 7 |
| 16 | Puszedli | APT | M | 17 | 24 | Y | <u>Y</u> | Y | 1090 | 7 |
| 17 | Csipesz | Mix | M | 32 | 24 | Y | <u>Y</u> | Y | 532 | 7 |

<sup>1</sup> Months. <sup>2</sup> Number of camera views recorded for each session. M<sup>1</sup> = Clip-on wireless microphone (Rode Wireless Go 2); M<sup>2</sup> = Sennheiser ME66 shotgun microphone; M<sup>3</sup> = Sennheiser ME64 cardioid microphone; F = Female; M = Male; Y = Yes; N = No; Breed abbreviations: BST = Belgian Shepherd Tervuren; NSCTR = Nova Scotia duck tolling retriever; APT = American Pit Bull Terrier. Recordings shown in bold and underlined indicate the recording used for annotation.

Dogs were recruited through social media announcements via the channels of the Department of Ethology at ELTE University, as well as through printed flyers. Inclusion criteria required dogs to be between 6 and 24 months of age, considered playful by their caregivers, and healthy at the time of participation. Both the caregiver and the dog were free to discontinue participation and leave the session at any time.

Participation was voluntary, and informed consent was obtained from all caregivers prior to testing. All procedures were approved by the institutional animal ethics committee in Hungary (approval number: ELTE-AWC-003/2026), in accordance with Good Scientific Practice guidelines and national legislation. The study complies with the ARRIVE guidelines^23^ and was carried out in accordance with the ASAB/ABS guidelines^24^.

## Data collection

Data were collected between March and May 2025 at the BARKS Lab, Department of Ethology, Eötvös Loránd University, Budapest, Hungary. The sessions took place at the Audio laboratory (5.46 m x 4.41 m). We used a camera system with seven Basler a2A 1920-51gcPRO IP cameras and a sound recording system consisting of a Zoom H4 Essential recorder, a clip-on wireless microphone: Rode Wireless Go 2; a Sennheiser ME66 shotgun microphone; a Sennheiser ME64 cardioid microphone; and two K6 power modules (Figure 1A).

**Figure 1.**
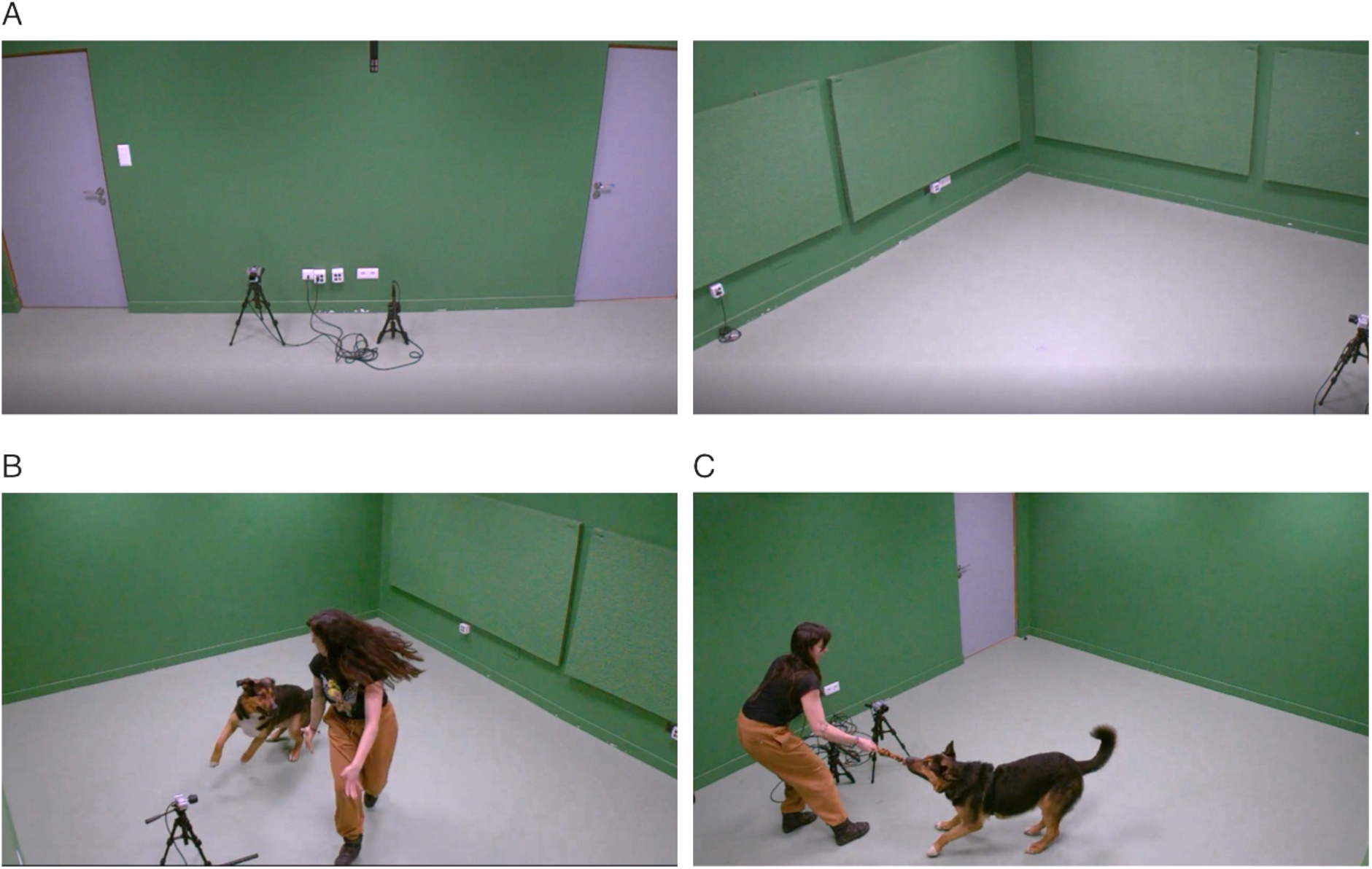
Experimental setup and examples of dog-human social play sessions. **(A)** Recording setup used during the experimental sessions. The left panel shows the view toward the entrance of the laboratory, including a camera used for close-up recordings and a Sennheiser ME66 shotgun microphone. The right panel shows a wider view of the laboratory. Examples of social play sessions recorded without toys **(B)** and with toys **(C)**, reflecting the typical play style of each dog-human dyad. Panels B and C show co-author Paula Pérez-Fraga interacting with Simon.

## Procedure

Before the recordings, caregivers completed a consent form and a brief survey providing information about their dogs, including age, breed, weight, and date of birth. As part of a pilot study, some dogs wore a Polar heart rate monitor, but the recordings were not usable, so heart rate data are not included in this dataset. Each play interaction lasted at least 3 minutes and continued until the dog was no longer engaged in play. Caregivers were instructed to interact with their dogs as they normally would during play and to try to elicit a playful and excited state. They were allowed to talk to their dogs and produce some noise; however, the aim was to obtain several audio segments without background noise and without toys in the dog’s mouth. To reduce potential recording noise, caregivers were asked to remove any metal dog tags or noisy collars before the session. Play sessions began with caregivers attempting to engage the dogs in play without making much noise or talking; if the dog did not become playful after approximately one minute, caregivers were then allowed to use more vocal encouragement. Similarly, sessions started without toys (Figure 1B), but if the dog was not willing to play after approximately 1–2 minutes, toys could be introduced (Figure 1C). A researcher (PP-F and/or CR) was present in the room during all sessions and occasionally provided guidance to caregivers without interfering with the dog-human social play.

## Data segmentation and annotations

From the raw recordings (Table 1), we selected segments in which dogs were actively playing with their caregivers. Portions of the recording containing researcher feedback, pauses, or other distractions were excluded so that each session corresponded to a continuous period of dog-human social play. In total, 30 sessions from 17 individual dogs were included in the dataset (1–3 sessions per dog). Session durations ranged from 34 to 613 seconds (mean = 249.40 seconds). The combined duration of the analysed recordings was 7482 seconds. When available (15 participants), the shotgun microphone recording was used for annotation due to its higher audio quality and lower perceived background noise (Table 1).

Dog sounds were annotated in the 30 dog-human social play sessions by three annotators: CR (7 sessions), Liene Kruskopa (2 sessions), and LP (21 sessions). Annotation was performed using spectrogram and waveform visualisation in Raven Lite Version 2.0.5^25^ to facilitate accurate identification of sound boundaries. Annotators followed a predefined annotation scheme to identify dog sounds in the recordings, marking the temporal boundaries (onset and offset) of each sound and assigning the corresponding label. The ethogram of sounds was compiled by PP-F and TF (Table 2). Importantly, sounds were not always annotated individually, but rather in bouts when two or more sounds of the same category occurred consecutively. The criterion for defining bouts was that no sound from different categories or noise could occur between sounds of the same category. This is especially important for panting, as almost all occurrences were annotated as bouts. Although not vocalizations, sneeze and shake events were also annotated. Sneezing has been described as a social signal in canids^26^ while body shaking has been suggested to function as a behavioural transition marker in dogs^27^. Additionally, although caregivers were instructed to produce as little noise as possible during the sessions, their vocalizations were annotated when present, as they form part of the dynamics of dog-human social play^8^.

**Table 2.**
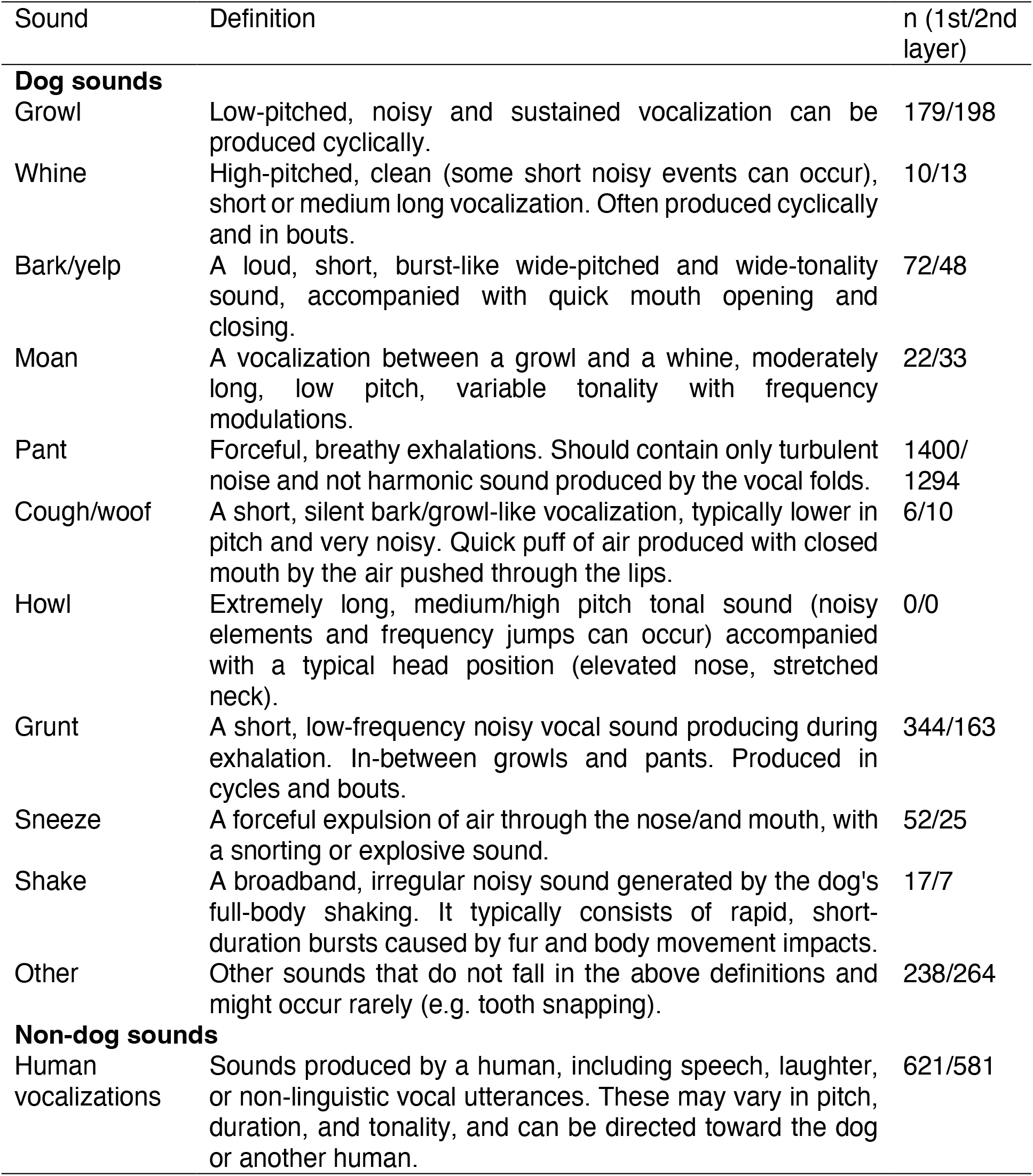
Sound category ethogram and annotation counts.

| Sound | Definition | n (1st/2nd layer) |
| --- | --- | --- |
| **Dog sounds** |  |  |
| Growl | Low-pitched, noisy and sustained vocalization can be produced cyclically. | 179/198 |
| Whine | High-pitched, clean (some short noisy events can occur), short or medium long vocalization. Often produced cyclically and in bouts. | 10/13 |
| Bark/yelp | A loud, short, burst-like wide-pitched and wide-tonality sound, accompanied with quick mouth opening and closing. | 72/48 |
| Moan | A vocalization between a growl and a whine, moderately long, low pitch, variable tonality with frequency modulations. | 22/33 |
| Pant | Forceful, breathy exhalations. Should contain only turbulent noise and not harmonic sound produced by the vocal folds. | 1400/1294 |
| Cough/woof | A short, silent bark/growl-like vocalization, typically lower in pitch and very noisy. Quick puff of air produced with closed mouth by the air pushed through the lips. | 6/10 |
| Howl | Extremely long, medium/high pitch tonal sound (noisy elements and frequency jumps can occur) accompanied with a typical head position (elevated nose, stretched neck). | 0/0 |
| Grunt | A short, low-frequency noisy vocal sound producing during exhalation. In-between growls and pants. Produced in cycles and bouts. | 344/163 |
| Sneeze | A forceful expulsion of air through the nose/and mouth, with a snorting or explosive sound. | 52/25 |
| Shake | A broadband, irregular noisy sound generated by the dog's full-body shaking. It typically consists of rapid, short-duration bursts caused by fur and body movement impacts. | 17/7 |
| Other | Other sounds that do not fall in the above definitions and might occur rarely (e.g. tooth snapping). | 238/264 |
| **Non-dog sounds** |  |  |
| Human vocalizations | Sounds produced by a human, including speech, laughter, or non-linguistic vocal utterances. These may vary in pitch, duration, and tonality, and can be directed toward the dog or another human. | 621/581 |

## Data records

The dataset described in this study is publicly available on Zenodo^28^. The dataset contains raw recordings, annotated play sessions, and metadata describing the participating dogs and recording conditions.

### 1. Samples of play sessions

Example video clips (10 seconds) are provided for each dog. These clips illustrate the dog-human social play interactions and provide a quick overview of the type of recordings and experimental conditions included in the dataset. The samples allow users to visually inspect the context in which sounds were produced and to become familiar with the structure and characteristics of the recordings.

### 2. Raw recordings

The raw data include audio and video recordings from 17 dogs collected during dog-human social play interactions at BARKS Lab, Budapest, Hungary. Video recordings capture the play sessions from multiple camera views. Audio was recorded using a Zoom H4 recorder with an external microphone, using two or three microphones depending on the recording setup. Recordings were saved in WAV format using PCM encoding, with a sampling rate of 48 kHz and a bit depth of either 16-bit or 32-bit. The raw recordings include the original camera recordings as well as synchronized videos with two angles of view and sound.

### 3. Play session recordings

Audio recordings corresponding to 30 dog-human social play sessions are included in the dataset as WAV files. Each audio file corresponds to one continuous play interaction between a dog and their caregiver. Most play session recordings are single-channel (mono) with a bit depth of 16-bit; however, sessions recorded with the Sennheiser cardioid microphone (P-05 and P-06) contain two channels (stereo), and P-01 was recorded at a bit depth of 32-bit. These technical details are documented in the metadata file.

### 4. Annotations

Dog sounds were annotated in the 30 play sessions using two annotation layers. The annotations provide the temporal boundaries (onset and offset) for each identified sound together with the assigned sound label. For the first layer, annotation files are tab-delimited text files exported from Raven Lite^25^ and contain the following fields: Selection (sequential annotation number), View (spectrogram view identifier); Channel (audio channel); Begin Time (s) (onset of the sound in seconds), End Time (s) (offset of the sound in seconds), and Annotation (a single letter label corresponding to the sound categories defined in Table 2: A = Growl, B = Whine, C = Bark/yelp, D = Moan, E = Pant, F = Cough/woof, G = Howl, H = Grunt, I = Other, J = Sneeze, K = Shake, L = Human vocalizations).

### 5. Metadata

It describes the sample and recording sessions in a tabular file (.csv) with semicolons as field delimiters. It includes information about dog identity, session number, recording date, and dog characteristics such as name, breed, sex, age, and weight. Additional fields document the recording equipment and technical audio parameters associated with each recording. Annotation files correspond directly to the audio recordings and reference the associated session identifiers, allowing users to align sound events with the original audio files.

### 6. Readme

This file provides an overview of the dataset structure, file organization, and naming conventions. It also includes guidance on how to navigate the dataset and links the different components (samples of the data, raw recordings, annotations, and metadata).

## Technical validation

After the initial annotation phase, a researcher with extensive experience in bioacoustics (PP-F) reviewed all annotation tables (n = 30) to ensure consistency and accuracy across sessions. To this end, all the text files from the first annotation layer generated in Raven were converted into Praat^29^ TextGrid files (Raven annotation start and end times were translated into interval boundaries, and labels translated into call type names, while human sound annotations were added as a separate interval tier), and the revision was carried out using a custom Praat^29^ script. During this the expert first checked and corrected the sound boundaries, then reviewed the labels and corrected both if necessary. This process results in an expert-reviewed version of the annotations for each session (Table 3). Both the initial annotations and the expert-reviewed annotations are included in the dataset as separate annotation layers. Table 2 shows the total annotation counts per sound category across both layers, and Table 3 shows the annotation counts per session across both layers.

**Table 3.** Descriptive data for play sessions (n = 30)

| Session | Dog ID | Duration (s) | Most common sound<sup>1</sup> |  | Total annotations (n)<sup>2</sup> |  |
| --- | --- | --- | --- | --- | --- | --- |
|  |  |  | First Layer | Second Layer | First Layer | Second Layer |
| 01 | 01 | 390 | Pant | Pant | 272 | 287 |
| 02 | 02 | 143 | Pant | Pant | 52 | 58 |
| 03 | 02 | 235 | Pant | Pant | 78 | 103 |
| 04 | 03 | 209 | Pant | Pant | 138 | 119 |
| 05 | 03 | 222 | Pant | Pant | 96 | 104 |
| 06 | 04 | 110 | Human voc. | Human voc. | 49 | 53 |
| 07 | 04 | 507 | Pant | Pant | 192 | 163 |
| 08 | 05 | 193 | Human voc. | Pant | 32 | 52 |
| 09 | 05 | 380 | Pant | Pant | 129 | 131 |
| 10 | 06 | 395 | Pant | Pant | 174 | 211 |
| 11 | 07 | 183 | Grunt | Pant | 113 | 190 |
| 12 | 07 | 95 | Human voc. | Human voc. | 64 | 53 |
| 13 | 07 | 248 | Pant | Pant | 117 | 89 |
| 14 | 08 | 408 | Pant | Pant | 111 | 85 |
| 15 | 09 | 199 | Pant | Pant | 49 | 62 |
| 16 | 09 | 166 | Pant | Pant | 48 | 40 |
| 17 | 10 | 148 | Grunt | Pant | 50 | 44 |
| 18 | 10 | 203 | Pant | Pant | 63 | 23 |
| 19 | 11 | 251 | Pant | Pant | 90 | 82 |
| 20 | 12 | 135 | Pant | Pant | 31 | 30 |
| 21 | 12 | 156 | Human voc. | Human voc. | 44 | 39 |
| 22 | 13 | 116 | Human voc. | Pant | 64 | 67 |
| 23 | 13 | 144 | Pant | Pant | 36 | 30 |
| 24 | 13 | 139 | Pant | Pant | 37 | 37 |
| 25 | 14 | 250 | Pant | Pant | 64 | 55 |
| 26 | 15 | 34 | Grunt | Pant | 14 | 13 |
| 27 | 15 | 178 | Grunt | Pant | 41 | 49 |
| 28 | 15 | 594 | Pant | Pant | 166 | 149 |
| 29 | 16 | 613 | Pant | Human voc. | 312 | 183 |
| 30 | 17 | 438 | Pant | Human voc. | 235 | 35 |

<sup>1</sup> Indicates the sound category with the highest frequency within each session.
<sup>2</sup> Total number of annotated events in each session.
Human voc. = Human vocalizations

Inter-annotator reliability was assessed by having an additional researcher with extensive experience in bioacoustics (TF) independently annotate 20% of the recordings using the same annotation ethogram (Table 2). These independent annotations were compared with the expert-reviewed annotations (textgrids). Agreement between annotators was evaluated both for the number of sounds detected and for the classification of sound types. Reliability in the number of sounds identified per recording was assessed using an intraclass correlation coefficient (ICC, two-way random-effects model), showing high agreement between annotations (ICC = 0.855, 95% CI = 0.703 – 0.932; F (25,25.3) = 12.4, p < 0.001). Agreement on sound category classification was evaluated using Gwet coefficient for each recording. The average coefficient was 0.786±0.112, indicating a good agreement between the expert raters.

Then, we conducted a classifier-based validation analysis to assess whether the annotated categories exhibit discriminable acoustic structure and to evaluate the internal consistency of the dataset. This analysis was performed separately for the first-layer annotations and the expert-reviewed second layer annotations. Acoustic features were extracted using the short-term feature extraction function of pyAudioAnalysis^30^ which computes frame-level descriptors over consecutive overlapping windows. Short-term features were computed using frame size of 0.010 seconds and a frame step of 0.005 seconds. For each annotated sound segment, the short-term feature sequence was extracted and summarized by computing the mean and standard deviation of each feature, yielding a fixed-length feature vector per segment. A linear support vector machine (SVM) classifier with a regularization parameter C = 1.0 was trained using a one-vs-rest multiclass scheme as implemented in scikit-learn^31^. Performance was quantified using macro-averaged F1 (macro-F1), which gives equal weight to each class and is therefore more appropriate than overall accuracy given the class imbalance across sound categories. Evaluation was performed using 5-fold grouped cross-validation, where groups corresponded to individual dogs, ensuring that samples from the same participants were never present in both training and test folds. Statistical significance was assessed using a permutation test on macro-F1 (N = 10,000). The complete cross-validation procedure was repeated after randomly shuffling class labels within each participant (thus preserving per-participant label composition), and the empirical null distribution of macro-F1 was used to compute the permutation p-value for the observed score. This procedure was applied independently to each annotation layer, yielding comparable performance metrics that reflect the internal consistency in each annotation set.

The linear SVM classification analysis yielded a macro-F1 of 0.241 ± 0.052 for the first layer annotations and 0.293 ± 0.024 for the expert-reviewed second-layer annotations (mean ± SD across grouped cross-validation folds). Permutation testing using 10,000 label-shuffled iterations (with labels shuffled within participants) showed that both observed macro-F1 scores were significantly higher than their empirical null distribution (p < 0.01, for both layers), supporting that the classifier captured category-relevant acoustic structure rather than being driven by individual-specific characteristics in each annotations set. Confusion matrices for both layers are shown in Figure 2.

**Figure 2.**
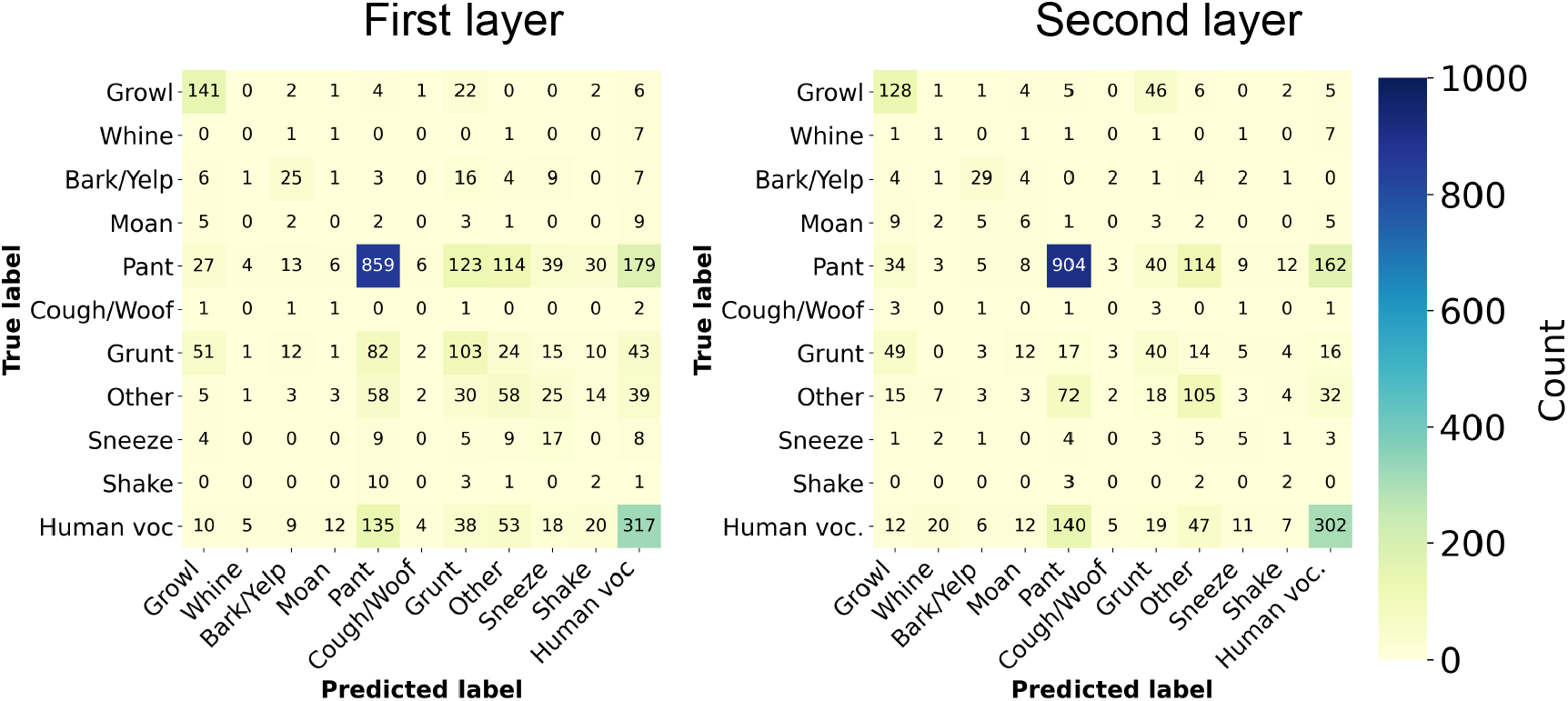
Confusion matrix of out-of-fold predictions from the linear SVM classifier for dog-human social play sounds. The matrix depicts classification patterns of the linear support vector machine (SVM) across annotated sound categories based on pooled out-of-fold predictions obtained during grouped cross-validation by participant for the first and second layer. Rows represent the true annotated labels, while columns represent the predicted labels. Diagonal elements indicate the number of correct predictions for each class. No sound was classified as howl. Human voc. = Human vocalizations

These results support the internal consistency of the annotated categories and the dataset’s suitability for reuse in comparative and machine-learning analyses of dog sounds during social play. However, the moderate classification performance suggests some acoustic overlap among categories, consistent with graded rather than strictly discrete vocal distinctions. Moreover, the validation relies on predefined acoustic features and a linear classifier, which may not capture all relevant structure present in the signals. Overall, the classifier-based validation results indicate that the annotated sound categories capture consistent acoustic structure that generalized across individuals, supporting the internal coherence of the dataset.

## Data availability

The dataset described in this paper is publicly available on the Zenodo repository (DOI: https://doi.org/10.5281/zenodo.18972388)

## Code availability

The code is available on: https://github.com/rhernandez00/bioacoustic-dataset

## Acknowledgements

We thank the dogs and their families and foster families for their time and enthusiasm in participating in this project. We are grateful to FAPF (https://www.fapfhungary.com), for allowing some of the dogs under their care to participate in this study. We would like to acknowledge Liene Kruskopa for her assistance with the annotations and data management, and Ernesto Zarco for his help in designing and disseminating the recruitment flyers, as well as for his assistance during data collection. We are also grateful to Sabrina Weinzettl and Emmanuelle Brogat for their administrative support. This research was enhanced by a collaboration at the Diverse Intelligences Summer Institute (DISI), funded by a grant from the John Templeton Foundation.

## Author contributions

**LVC**: Conceptualization, Methodology, Formal analysis, Data Curation, Writing - Original Draft, Writing - Review & Editing, Visualization, Supervision, Project Administration, Funding acquisition. **PP-F**: Conceptualization, Methodology, Investigation, Validation, Writing - Review & Editing, Project Administration, Funding acquisition. **RH-P**: Software, Validation, Formal analysis, Data Curation, Writing - Review & Editing, Visualization. **LP:** Investigation, Data Curation, Writing - Review & Editing. **IW**: Data Curation, Writing - Review & Editing. **CR**: Investigation, Writing - Review & Editing. **TF**: Methodology, Validation, Resources, Writing - Review & Editing. **SW**: Conceptualization, Writing - Review & Editing, Funding acquisition. **LH**: Resources, Writing - Review & Editing, Supervision. **CL**: Resources, Writing - Review & Editing, Supervision.

## Competing interest

The authors declare no competing interests.

## Funding

This publication was made possible through the support of Grant 63138 from the John Templeton Foundation. The opinions expressed in this publication are those of the author(s) and do not necessarily reflect the views of the John Templeton Foundation. This research was funded in whole or in part by the Austrian Science Fund (FWF): 10.55776/ESP602 and by the European Research Council (ERC) under the European Union’s Horizon Europe research and innovation programme (101125731). PPF was supported by the EKÖP-25-4-II University Excellence Scholarship Program by the National Research, Development and Innovation Office (EKÖP-25-4-II-ELTE-900). For open access purposes, the author has applied a CC BY public copyright license to any author-accepted manuscript version arising from this submission.

## Notes

https://doi.org/10.5281/zenodo.18972388

### Competing Interest Statement

The authors have declared no competing interest.

